# Reconstructing the pressure field around a swimming fish using a physics-informed neural network

**DOI:** 10.1101/2023.02.27.530217

**Authors:** Michael A. Calicchia, Rajat Mittal, Jung-Hee Seo, Rui Ni

## Abstract

Hydrodynamic pressure is a physical quantity that is utilized by fish and many other aquatic animals to generate thrust and sense the surrounding environment. To advance our understanding of how fish react to unsteady flows, it is necessary to intercept the pressure signals sensed by their lateral line system. In this study, the authors propose a new, non-invasive method for reconstructing the instantaneous pressure field around a swimming fish from 2D particle image velocimetry (PIV) measurements. The method uses a physics-informed neural network (PINN) to predict an optimized solution for the velocity and pressure fields that satisfy in an ℒ_2_ sense both the Navier Stokes equations and the constraints put forward by the measurements. The method was validated using a direct numerical simulation of a swimming mackerel, *Scomber scombrus*, and was applied to empirically obtained data of a turning zebrafish, *Danio rerio*. The results demonstrate that when compared to traditional methods that rely on directly integrating the pressure gradient field, the PINN is less sensitive to the spatio-temporal resolution of the velocity field measurements and provides a more accurate pressure reconstruction, particularly on the surface of the body.

## Introduction

Aerodynamic or hydrodynamic pressure is a physical quantity that is utilized by animals to both generate thrust (Akhtar et al., 2007; Zhang et al., 2015; Anderson et al., 2017; Dagenais et al., 2020; Thandiackal et al., 2020; Tack et al., 2021; Saadat et al., 2021; Han et al., 2022) and sense the surrounding environment (Liao et al., 2003; Liao, 2006; Ristroph et al., 2008; McHenry et al., 2009; Ashraf et al., 2017; Verma et al., 2018; Halsey et al., 2018; Li et al., 2020). For example, fish have a sensory system, i.e., the lateral line, for detecting the rapidly changing pressure of the flow (Ristroph et al., 2020). To understand how fish react to unsteady flows, it is necessary to intercept the pressure signals received by the fish; however, it is challenging to do this instantaneously in a non-invasive manner.

The most utilized non-invasive method is to reconstruct the pressure field from velocity measurements (van Oudheusden., 2013). Traditionally, there have been two main categories of this approach. The first computes the pressure field from the Poisson equation, i.e., as shown below for an inviscid flow (Fujisawa et al., 2005; de Kat et al., 2012; Shams et al., 2015; Neeteson et al., 2015; Pirnia et al., 2020).

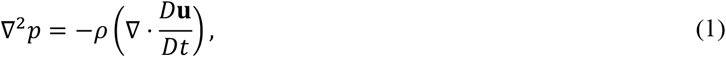

where *p* is the pressure, **u** is the velocity vector, *ρ* is the fluid density, and *D*/*Dt* is the material derivative. However, Charonko et al. (2010) and Pan et al. (2016) have shown that the Poisson-based solvers are sensitive to the grid resolution, flow type, velocity measurement errors, the shape of the immersed body, and the type of boundary conditions that are applied. Furthermore, as Dabiri et al. (2014) suggested, when applied to the study of animal locomotion under low or moderate Reynolds number (*Re*), it is difficult to predetermine the appropriate boundary condition at the fluid-body interface. Therefore, the pressure reconstruction could benefit from new methods that are less sensitive to these constraints.

The second category of techniques for pressure reconstruction is the direct integration of the pressure gradient along multiple different paths (Liu et al., 2006; 2013; Dabiri et al., 2014; Liu et al., 2021; Agarwal et al., 2021), as shown below.

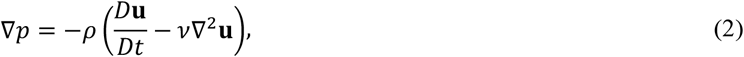

Here, *ν* is the kinematic viscosity of the fluid. Multi-directional integration schemes utilize the scalar property of pressure, i.e., its local value is independent of the path taken, to improve the accuracy of the pressure estimation. Using this approach, Dabiri et al. (2014) developed an unsteady pressure reconstruction algorithm, known as Queen 2.0, to study animal locomotion (Dabiri et al., 2020; Dagenais et al., 2020; Siala et al., 2020; Costello et al., 2021; Gemmell et al., 2021; Kasoju et al., 2021; Thandiackal et al., 2021; Guo et al., 2022).

However, Queen 2.0 does have its limitations. Firstly, to integrate the pressure gradient, a zero-pressure boundary condition is applied at all external boundaries, which is not always accurate. As demonstrated by He et al. (2020), when, for example, the wake of a turbulent jet crosses one of the boundaries, the pressure reconstruction by Queen 2.0 becomes less accurate. Secondly, Queen 2.0 and most of the direct integration methods, do not incorporate information at the fluid-body interface into the pressure reconstruction. This is typically done because the velocity measurements, especially those obtained in typical PIV experiments, nearest to the body are often unreliable. Thus, to avoid this error from propagating to the pressure estimation, the integration paths are terminated before reaching the fluid-body interface. To then obtain the surface pressure, one would typically have to extrapolate from the nearest neighbor node in the surrounding pressure field. Pirnia et al. (2020) demonstrated that such an approach can provide a very accurate prediction of the surface pressure around stationary objects. However, the error increases greatly when the object is free to deform. They also showed that by incorporating the kinematics of the immersed body into the pressure reconstruction algorithm, the relative error in the surface pressure prediction can be sufficiently reduced.

These results stress the need to have a pressure reconstruction algorithm that: 1) provides the user with the flexibility to alter the applied boundary conditions and 2) incorporates the kinematics of the undulating body into the pressure reconstruction. In recent years, new types of pressure reconstruction algorithms have been developed (Wang et al., 2017; Jeon et al., 2018; Huhn et al., 2016; Cai et al., 2020; Wang et al., 2018; He et al., 2020); although these methods have made significant progress in other aspects of pressure reconstruction from velocity measurements, they do not correct the highlighted limitations of Queen 2.0. Furthermore, their applicability to flow fields involving actively deforming bodies remains relatively untested.

Therefore, in this paper, we propose a new method to reconstruct the pressure field around undulating bodies based on physics-informed neural networks (PINNs) (Cai et al., 2022). The most important benefit of using PINNs is their flexibility. PINNs can deal with any boundary condition or no boundary condition; they do not need to deal with the complex grid designs required to incorporate the kinematics of the immersed body; they are less sensitive to the spatio-temporal resolution and noise, and they can patch the results in regions where velocity field data is not available (Cai et al., 2021, Di Leoni et al., 2022, Du et al., 2023).

Previous research performed by Raissi et al. (2020) has utilized PINNs to reconstruct the pressure field around a stationary object. To build on this work, the authors will apply the method to reconstruct not only the pressure field around a swimming fish but also the pressure signals sensed by its lateral line.

## Materials and methods

### Physics-informed neural networks

The general idea of the proposed method is not to derive pressure from the velocity field via integration, but to seek an optimized solution that simultaneously satisfies the governing equations and the constraints put forward by the measurements in an ℒ_2_ sense. The machine learning architecture provides an efficient way to meet these two requirements by iteratively updating the trainable parameters of the network to minimize a loss function, ℒ. The loss function can be decomposed into four main terms: the measured data (ℒ_*data*_), the imposed initial conditions (ℒ_*𝒥𝒞*_), the imposed boundary conditions (ℒ_*𝔅𝒞*_), and the governing equations (ℒ_*𝒩𝒮*_). Thus, the loss function can be expressed as:

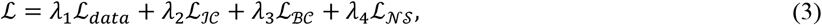

where *λ*_1−4_ are the weighting coefficients for the different loss terms. In this study, a fully connected feed-forward neural network is used to approximate the solution of the Navier-Stokes equations to recover the two-dimensional pressure field around a swimming fish. The PINN takes the spatio-temporal coordinates as inputs and performs a series of algebraic operations as they pass through twelve hidden layers, each of which contains 120 neurons. The output of the last layer, *K*, is used to approximate the solution of the Navier-Stokes equations. If the input variables to the *k*^*th*^ hidden layer are denoted **z**^*k*^ (k=1,2, 3,…K-1), then the neural network can be represented as

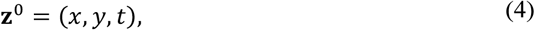

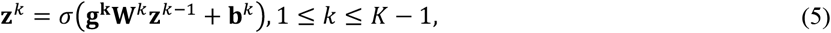

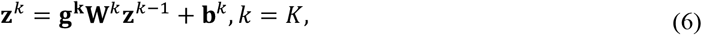

where *x* and y denote the spatial coordinates, *t* denotes the temporal coordinates, **W**^*k*^, **b**^**k**^, and **g**^**k**^ denote the trainable parameters of the network: weights matrix, bias and gamma vectors, respectively, and σ(·) denotes the activation function. In this study, a sigmoid activation function was used. To determine an appropriate network size, a parametric study was performed in which the number of layers and neurons per layer were systematically varied. For each network size the global relative root mean square error in the velocity and pressure fields were computed. In the supplementary material, Fig. S1 shows that a network size of twelve layers consisting of 120 neurons provided the most accurate solution for the pressure field.

In this application, how accurately the PINN predictions match the measured time-series of the two-dimensional velocity fields can be quantified by the following data loss term.

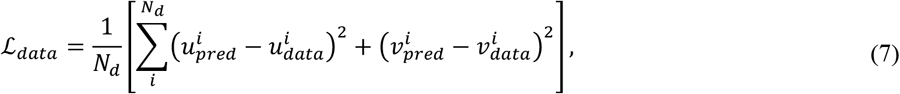

where *N*_*d*_ is the number of training data points sampled at each iteration, and *u* and *ν* are the lateral and transverse velocities, respectively. The subscript “*pred*” refers to the predictions by the PINN, and the subscript “*data*” refers to the velocities obtained from the simulation or PIV results. The training data includes the velocity vectors in the domain over the entire time.

To enforce the physics of the problem, the residuals of the Navier-Stokes equations are also evaluated. In general, the equation loss term consists of the residuals of the dimensionless momentum equations and continuity equation. However, since a two-dimensional slice is extracted from a three-dimensional velocity field, the divergence free condition is not enforced. Furthermore, since the third component of the velocity field is missing, the product of the out-of-plane velocity and the spatial derivative of *u* and *ν* in that direction is assumed to be negligible. Therefore, it is important to stress that the current method is only applicable to cases where three-dimensional effects are weaker. This can be achieved by ensuring the PIV plane passes through the midline of the fish’s body and that the fish’s motion lies within this plane. The Navier-Stokes residuals utilized in this framework are shown as follows:

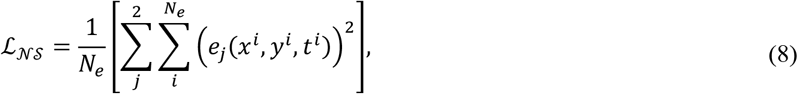

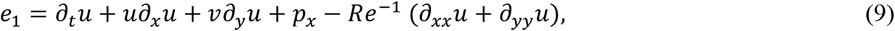

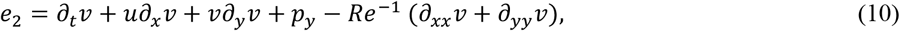

Here, *N*_*e*_ is the number of data points sampled at each iteration to evaluate the Navier-Stokes residuals. The partial derivatives in the governing equations are computed using automatic differentiation (Baydin et al., 2018), which calculates the derivatives of the outputs (*u, ν, p*) with respect to the network inputs (*x, y, t*) directly in the computational graph, without any finite differencing methods utilized in more classical computational methods. It is important to stress that the Navier-Stokes residuals can be evaluated at points where measured data is not available thus providing a means for increasing the resolution of the measured data, which is grounded in physics.

The applied initial and boundary conditions depend on the problem. Since the network is in essence solving the Navier-Stokes equations, the applied initial and boundary conditions can involve either pressure or velocity. Compared with other methods relying on integrating pressure from boundaries to the point of interest, this method does not require *a priori* knowledge of the boundary conditions and certainly does not enforce the wrong boundary condition when it is not available. Furthermore, PINNs do not rely on a traditional Cartesian grid because they simply take any spatio-temporal coordinate as inputs and outputs velocities and pressure. This feature is extremely helpful in dealing with complex animal locomotion problems because the undulating body and the fluids grid do not always coincide with one another. But for PINNs, there is no need to extrapolate from a grid to the body or back. The kinematics of the body can be input as a boundary condition into the network with ease.

For all cases in this study, a non-penetration boundary condition is enforced on the surface of the fish’s body, through which information of the fish kinematics is utilized. Therefore, the boundary condition is a measure of how well the PINN prediction matches the measured velocity normal to the fish’s body. In addition, boundary conditions can be enforced at external boundaries. These may include a zero-pressure boundary condition and an inlet velocity boundary condition. The boundary condition loss terms that were enforced in this study are shown as follows.

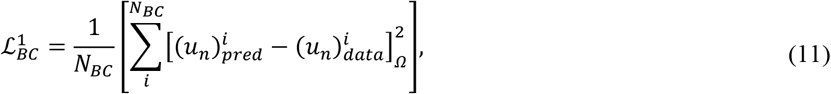

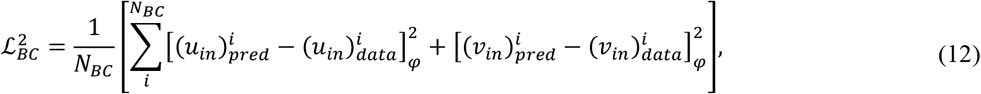

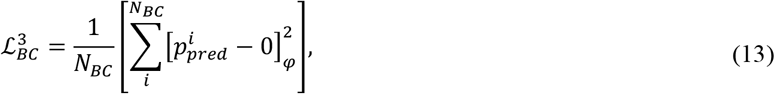

where *u*_*n*_ denotes the normal velocity, *u*_*in*_ denotes the lateral component of the inlet velocity, *ν*_*in*_ denotes the transverse component of the inlet velocity, *Ω* denotes the spatio-temporal coordinates of the fish’s body, *φ* denotes the spatio-temporal coordinates at the domain boundaries, and *N*_*BC*_ denotes the number of points on the boundary that were sampled at each iteration. There were no initial conditions applied to any of the cases in this study.

To minimize the loss function and optimize the trainable parameters of the network, the ADAM optimizer was used (Kingma et al., 2015). The mini-batch size was set to 10,000. Therefore, at each iteration, a maximum of 10,000 spatio-temporal points were randomly sampled from the entire training dataset to evaluate the terms of the loss functions. The PINN was trained on a NVIDIA a100 graphics card. For each case studied, the network was trained for 1500 epochs, or 1500 passes through the entire dataset, and took approximately ten hours to complete. As shown in the supplementary material, 1500 epochs sufficiently balances the accuracy of the PINN predictions with the computational cost.

It is important to note that the purpose of the PINN framework is to uncover hidden information from visualizations of the flow field. For this application, the goal is to recover pressure from velocity measurements. Therefore, for every new velocity field, the network must be retrained to obtain the corresponding pressure field. The trained network is not meant to predict the pressure field for a wide range of different flow types nor is it meant to be used to develop reduced order models.

In theory, the PINN method can be applied to study undulatory locomotion over a range of Reynolds numbers if the animal’s oscillatory motion is primarily two-dimensional and lies in the same plane as the PIV data. To satisfy these two conditions, two datasets of the flow produced by the oscillatory motion of carangiform swimmers were selected. One is the direct numerical simulation (DNS) of a swimming mackerel, which will be used to quantify the accuracy of the method. The other one is an experimental dataset of a turning zebrafish, *Danio rerio*.

### Single fish validation dataset

To test the accuracy of the proposed method, it was applied to a direct numerical simulation of a swimming fish using the ViCar3D, a sharp-interface immersed boundary method (Mittal et al., 2008). The 3D model of the fish is based on the common Mackerel (*Scomber scombrus*). The fish model consists of body and caudal fin, and the caudal fin is modelled as a zero-thickness membrane. A carangiform swimming motion is prescribed by imposing the lateral displacement of the fish body and fin using the following prescription: *Δy*/*L* = *A*(*x*)*sin*(*kx* − 2*πft* + *ϕ*) ; *A*(*x*) = *a*_0_ + *a*_1_(*x*/*L*) + *a*_2_(*x*/*L*)^2^ where *Δy* is the lateral displacement, *L* is the body length, *x* is the axial coordinate along the body starting from the nose, *f* is the tail beat frequency, *ϕ* is a phase, and *A*(*x*) is the amplitude modulation function. The parameters are set based on literature (Videler et al., 1987) to the following values: *a*_0_ = 0.02, *a*_1_=-0.08, and *a*_2_=0.16. The wave number is set to *k* = 2*π*/*L* and the flow Reynolds number based on the body length and tail beat frequency, Re_*L*_ = *L*^2^*f*/*ν*, is set to 5,000.

In the present simulation, the swimming motion is imposed on a ‘tethered’ fish and a flow velocity is prescribed at the inflow boundary, such that the net force on the fish is nearly zero, thereby simulating self-propelled swimming with net zero acceleration. The fish body and caudal fin are meshed with triangular surface elements and immersed into the Cartesian volume mesh which covers the flow domain. The flow domain size is set to 8*L* × 10*L* × 10*L* and this is discretized on a very dense grid with 640 × 320 × 240 (about 49 million) Cartesian cells. The minimum grid spacing (cell size) is 0.005*L* and the body length is covered by 200 grid points. The time-step size used in the simulation is *Δt* = 0.001/*f*, which resolves one tail beat cycle with 1000 time-steps. A no-slip, no-penetration boundary condition is applied on the moving fish body and fin surfaces by using the sharp-interface, immersed boundary method and a zero-gradient velocity boundary condition is applied on the other domain boundaries except the inflow. For the pressure, a zero gradient boundary condition is applied on the fish body as well as all the outer boundaries (Seo et al., 2022).

A 2D slice at a z-plane cutting through the midline of the fish’s body was extracted from the 3D velocity field. The 2D velocity and pressure gradients on this plane provide the hydrodynamic signals that a fish would sense with its lateral line. In addition, the simulation results fully define the fish’s motion as the velocities at multiple points along the fish’s body are known. The data extracted from the simulation results, which contains a time series of velocity fields and defined fish kinematics, were meant to replicate a dataset that can be obtained through experiments. For example, the velocity field on a plane cutting through the midline of a fish’s body can be obtained through particle image velocimetry (PIV), and the fish kinematics could be captured by imaging the silhouette of the fish body. The spatio-temporal resolution of the 2D velocity field on this DNS slice was made coarser to replicate data that would be obtained from a PIV experiment with either a large field of view or insufficient resolution.

To test the sensitivity of the proposed method to the spatial resolution, the PINN was trained using velocity fields with a grid size of 0.02*L*, 0.04*L*, 0.06*L*, 0.08*L*, and 0.1*L*, which would respectfully consist of 50, 25, 17, 13, and 10 grid points along the length of the fish’s body. For this study, a temporal resolution of 0.02*T* was used. Here, *L* is the body length of the fish, and *T* is the time corresponding to the fish’s motion, i.e., one period of the tail beating motion. To test the sensitivity of the proposed method to the temporal resolution, the PINN was trained using velocity fields with a time step of 0.02*T*, 0.04*T*, 0.06*T*, 0.08*T*, and 0.1*T*. For this study a spatial resolution of 0.02*L* was used. As previously mentioned, the Navier-Stokes residuals shown in Eqns 8-10 can be evaluated at any points in the domain and not necessarily at points where measurement data are available. Therefore, in the spatio-temporal resolution study, the Navier-Stokes residuals were always evaluated on the finest grid even as the velocity measurements became coarser.

To evaluate the accuracy of the pressure reconstruction along the surface of the body across all time steps for each spatio-temporal resolution tested, the relative global root mean square error (RMSE) was computed as follows.

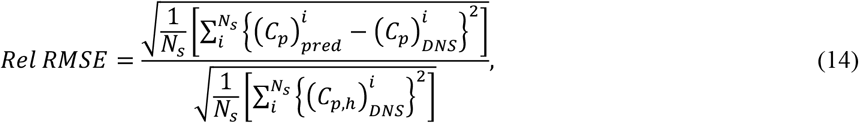

where *N*_*s*_ is the number of surface points, and 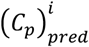 and 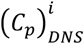 represent the non-dimensional surface pressure predicted by either the PINN or Queen 2.0 and the simulation data, respectively. 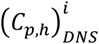 represents the non-dimensional pressure at the fish’s head. The authors chose to normalize the global RMSE by the head pressure because the surface pressure for most of the body is close to zero. Furthermore, all data points on the body located at *x* > 0.9*L* were excluded from the error calculation. As shown by Fig. S3 in the supplementary material, in this region the flow becomes highly three-dimensional and thus naturally the errors in the pressure field will be much larger.

In addition to the resolution of the velocity measurements, it is also expected that the level of noise in the velocity data will affect the accuracy of the PINN’s prediction. To test the sensitivity of the proposed method to noise, various levels of artificial white noise were systematically added to the velocity data on the 2D plane extracted from the DNS dataset. The details of this study and its results are available in the supplementary material.

For each dataset analyzed, the loss function to be minimized consisted of the two-dimensional velocity data loss terms, the residuals of the x and y momentum equations, and the boundary condition loss terms. To incorporate the kinematics of the fish’s body, a non-penetration boundary condition was applied on the surface of the fish’s body. In addition, a zero-pressure boundary condition was applied at the top and bottom boundaries since they were considerably far away from the fish’s motion. An inlet velocity boundary condition was also applied. However, no boundary condition was added to the outlet where the pressure is significantly affected by the wake. Lastly, the divergence free condition was not enforced since a two-dimensional slice was extracted from a three-dimensional velocity field.

For this application, *λ*_1_ and *λ*_3_ in Eqn 3 were set to 100, λ_2_ was set to zero, and *λ*_4_ was set to unity. As shown in the supplementary material, Fig. S2 demonstrates that by applying these weights, the PINN can more accurately recover the pressure field and provide a better prediction of the surface pressure. The choice of weights is consistent with that reported by Cai et al. (2021).

### Empirical dataset

To test the proposed method on empirical velocity field data, the PINN was used to reconstruct the two-dimensional pressure field around a turning zebrafish, *Danio rerio*. The velocity fields were obtained from PIV experiments performed by Thandiackal et al. (2020).

Before implementing the PINN, the velocity field grid points inside the fish’s body were removed from the dataset. Then, the velocity in the direction normal to the zebrafish’s body was computed at all time steps. Lastly, since the PINN utilizes the non-dimensional form of the Navier-Stokes equations, the spatio-temporal coordinates and the velocities were non-dimensionalized. The characteristic time was the turning time (0.15 s), the characteristic length was the zebrafish body length (22 mm), and the characteristic velocity was computed by dividing the center of mass displacement by the turning time. The Reynolds number for this case was 918. The dimensionless grid had a spatial resolution of 0.02 and a temporal resolution of 0.03. A total of 36-time steps were included in the training dataset.

For the empirical dataset, the loss function to be minimized consisted of the two-dimensional velocity data loss terms, the x and y momentum equations, and a loss term needed to enforce a non-penetration boundary condition on the surface of the fish’s body. In this case, the zero pressure boundary conditions were not enforced at any of the boundaries because the zebrafish is relatively close to the top and left boundaries at various points throughout its turning motion. It is uncertain whether the pressure field induced by the fish motion would affect the boundaries. When such uncertainty exists, it is better not to enforce the zero-pressure boundary condition; rather, one should allow the PINN to learn what the pressure at the boundaries should be based on all the information provided during the training process. The same weighting factors used for the simulation data were applied in this experimental data as well.

## Results and Discussion

Fig. 1 compares the instantaneous pressure field at *t* = 0.2*T* obtained from the simulation to that predicted by the PINN and Queen 2.0. The results were computed from the simulation dataset with a spatial resolution of 0.02*L* and a temporal resolution of 0.02*T*.

**Fig. 1.**
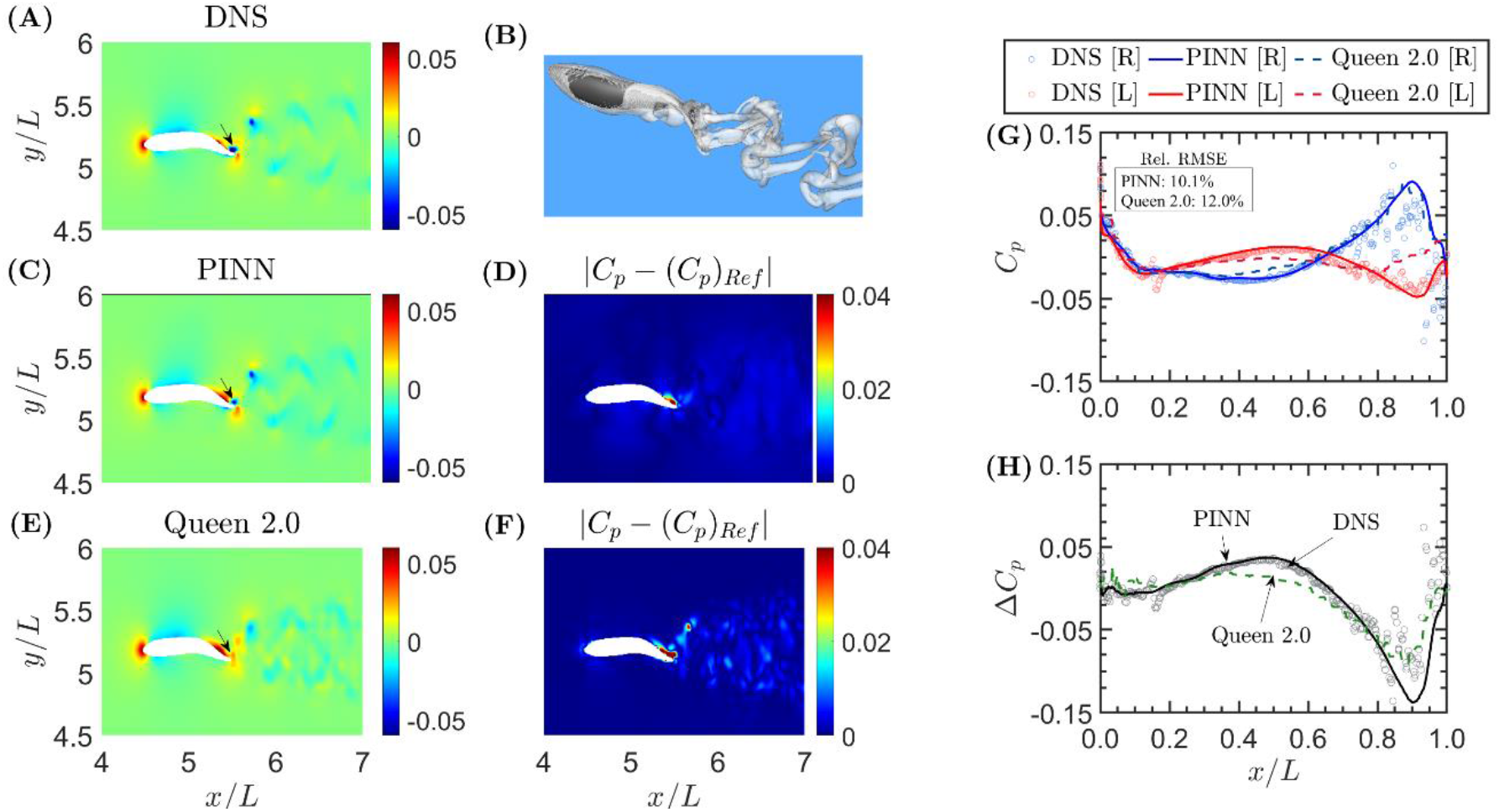
Comparison of the 2D pressure field and surface pressure profiles predicted by PINN and Queen 2.0. (A) the 2D pressure data cut through the mid-plane of the fish from (B) the full 3D DNS results. (C)-(F) the reconstructed pressure fields and their associated absolute error for each method. The arrows denote a region where the localized error in the Queen 2.0 prediction is much larger than PINN. (G) the surface pressure profiles, including the DNS results (circles) and the prediction from the PINN (solid) and Queen 2.0 (dashed), on the right (blue) and left (red) sides of the fish body. (H) the pressure difference between the left and right side of the body predicted by each method.

The PINN effectively captures the high-pressure region near the head, the large pressure variation near the tail, and the pressure fluctuation in the wake. In comparison, Queen 2.0 captures the high-pressure region at the head but is less accurate near the tail. In fact, Queen 2.0 does not capture the negative pressure region on the right side of the fish’s tail (pointed by the arrow in Fig. 1A,C,E) and instead predicts a region of positive pressure. This results in localized errors that are much larger than those obtained by the PINN.

### Results for validation dataset

The decrease in the errors predicted by the PINN in the tail region can be attributed to the fact that it utilizes the fish’s kinematics as another constraint in the pressure reconstruction and resolves the pressure field up to the fluid-body interface, whereas Queen 2.0 does not. Furthermore, Queen 2.0 enforces a zero-pressure boundary condition on the right side of the domain. This is not an accurate boundary condition since the vortices shed by the beating tail pass through the boundary and results in a non-zero pressure. For the Queen 2.0 reconstruction, in the wake region certain areas exhibit higher errors than others. This is most likely because in these localized regions the multi-directional integration scheme is not able to sufficiently mitigate the error introduced by applying a zero-pressure boundary condition on the right side of the domain. For PINNs, there is no need to enforce this boundary condition and introduce the associated error into the pressure reconstruction. Thus, for the PINN reconstruction, the error in the wake region is more uniform. The results in Fig. 1 demonstrate how the PINN can overcome limitations of Queen 2.0 and provide an accurate reconstruction of the pressure field surrounding a swimming fish.

It is important to note that both algorithms produced an increased error in regions where the out-of-plane velocities are non-negligible (i.e., in the tail and wake region). This is unsurprising since only the 2D flow field was used during the pressure reconstruction. A more detailed discussion of this result can be found in the supplementary material.

To better understand the signals that a fish is sensing with its lateral line, the pressure reconstruction method must accurately predict the pressure on the surface of the fish’s body. To obtain the surface pressure using Queen 2.0, one must extrapolate from the reconstructed pressure field. As was done in Thandiackal et al. (2020), the pressure at a point on the body is typically assumed to be the pressure at the closest grid point. In contrast, the PINN provides the ability to predict the surface pressure directly without any need for extrapolation.

Fig. 1G,H compares the surface pressure on the left (red) and right (blue) side of the fish’s body and the pressure difference between the two sides as predicted by the PINN and Queen 2.0 to that obtained from the simulation. The pressure difference profiles are included because Ristroph et al. (2020) have suggested that the pressure difference is a quantity that fish can sense. The results demonstrate that, for most of the sensing region of the fish, both methods can accurately predict the surface pressure with the PINN being slightly more accurate particularly in the tail region. This can be confirmed quantitatively by computing the relative RMSE in the surface pressure according to Eqn 14. The PINN has a relative error of 10.1% whereas Queen 2.0 has an error of 12.0% at this time step.

### Results for the spatio-temporal resolution study

The benefit of using PINNs becomes more apparent as the spatio-temperal resolution of the velocity field deteriorates. Fig. 2 compares the instantaneous surface pressure at *t* = 0.3*T* (top row) and *t* = 0.6*T* (bottom row) on the right side of the fish’s body predicted by the simulation to that predicted by the PINN and Queen 2.0 for each spatial and temporal resolution tested. For most of the body, the pressure predictions by the PINN collapse onto the simulation results for each of the spatio-temporal resolutions tested. Although at the coarsest resolution tested, comparatively larger deviations can be observed in the tail region. This result indicates that, for most of the body, the accuracy of the pressure reconstructed by the PINN is not that sensitive to the spatio-temporal resolution of the measured velocity field. For Queen 2.0, the accuracy of the surface pressure prediction greatly decreases as the resolution becomes coarser. The coarser the PIV grid, the farther the distance between the fish surface to the nearest PIV grid point. Since Queen 2.0 requires extrapolation from the grid to the surface, a longer extrapolation distance results in a larger error, as expected. The decrease in accuracy as a function of the temporal resolution, on the other hand, can most likely be attributed to the fact that the finite difference approximation of the material derivative shown in Eqn 2 becomes less accurate with a larger time step.

**Fig. 2.**
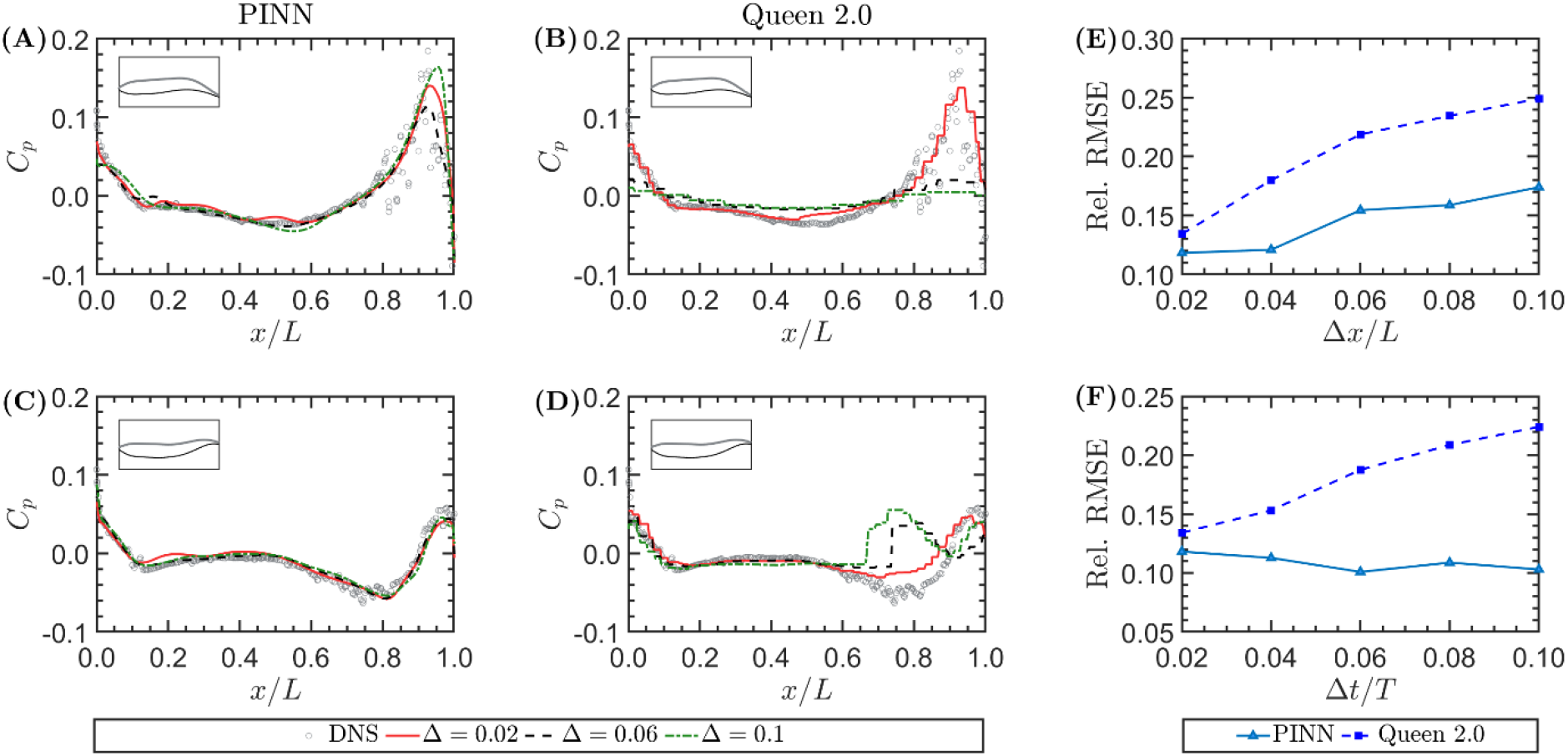
Surface pressure profiles reconstructed from the velocity data using PINN and Queen 2.0 at different (A)-(B) spatial and (C)-(D) temporal resolutions: 0.02 (red), 0.06 (blue), and 0.1 (green) for two different times. (E)-(F) The relative global root mean square error in the surface pressure prediction obtained by both methods as a function of (E) the spatial and (F) temporal resolution of the PIV data.

Fig. 2E,F shows how the relative global RMSE in the surface pressure prediction by each method varies as a function of the spatial and temporal resolution. For Queen 2.0, the error quickly grows as the resolution becomes coarser, but for the PINN the error profile remains relatively flat across all temporal resolutions tested. The error only begins to really rise as the spatial resolution exceeds 0.04*L*. Note that the errors reported in Fig. 2E,F only represent the global average, the improvement of local pressure prediction could be much larger than these values.

### Results for empirical dataset

Fig. 3A-F compares the instantaneous velocity field obtained from the PIV experiments to that reconstructed by the PINN. The PINN can accurately reconstruct the velocity fields, with absolute errors not exceeding 0.1. Since the optimization process was regularized by the Navier-Stokes residuals, an accurate reconstruction of the velocity field would imply that the reconstructed pressure field is also accurate. Fig. 3G,H shows the reconstructed pressure field around a turning zebrafish and the pressure distribution along its body directly predicted by the PINN. Although it is difficult to make a direct comparison to Queen 2.0 since the ground truth pressure field is unknown, the results from the validation case suggest that the PINN prediction would be more accurate because it resolves the pressure field all the way to the body, it incorporates the zebrafish’s kinematics into the pressure reconstruction, and does not enforce a zero pressure boundary condition since the zebrafish’s motion may induce flows crossing the boundaries.

**Fig. 3.**
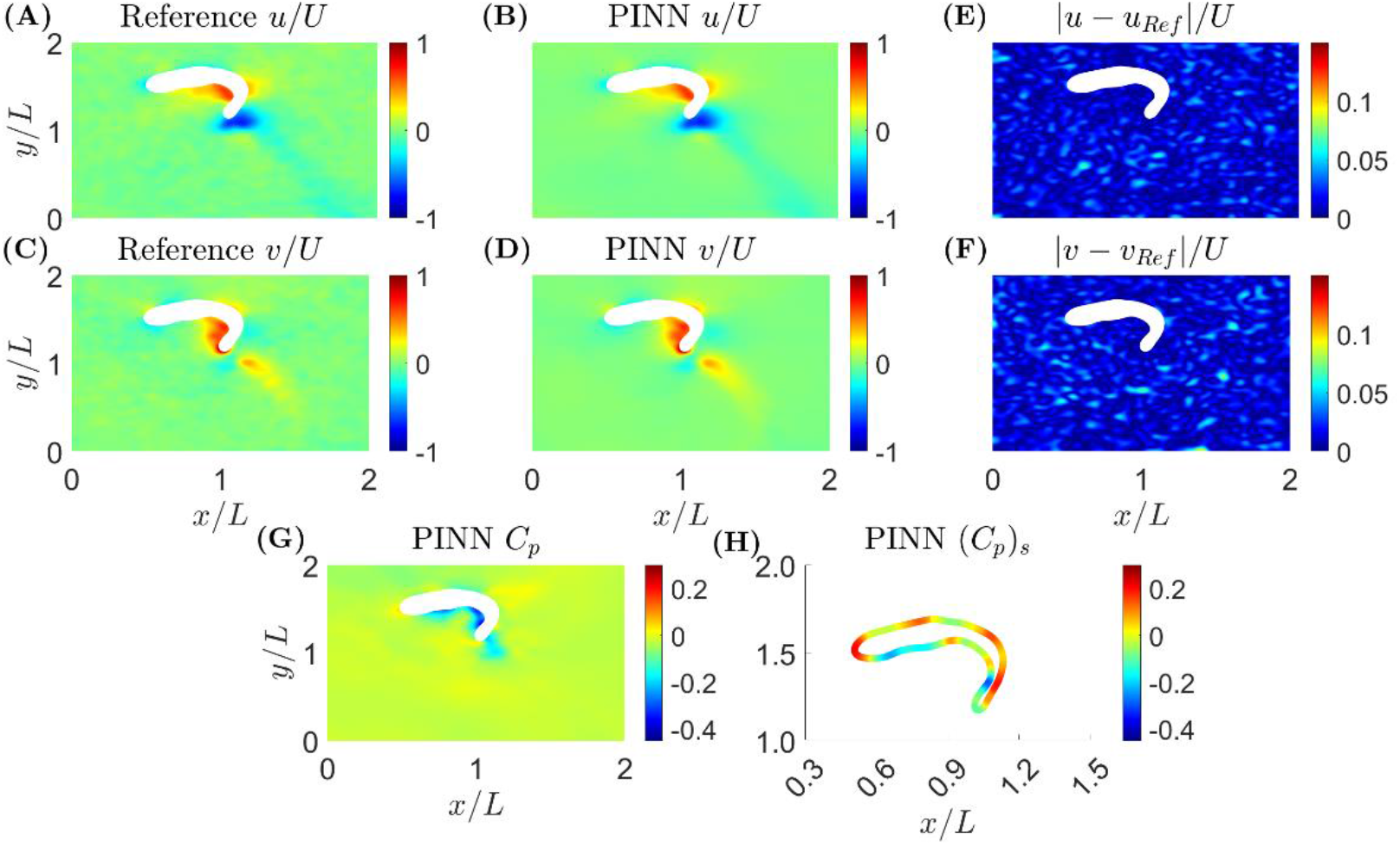
Reconstructed velocity and pressure fields for empirical dataset. (A)-(F) The predicted velocity fields versus the direct PIV measurements and their differences. (G)-(H) the instantaneous pressure field and surface pressure predicted by the PINN

### Comparative advantage

In this paper, a machine learning-based method for reconstructing the pressure field around an undulating body from 2D PIV data was developed. PINNs provide the user with greater flexibility in applying boundary conditions and provide a framework for incorporating the kinematics of the undulating body into the pressure reconstruction process without requiring any deforming grids. When compared to Queen 2.0, at the highest resolution of the PIV data, PINNs provide a small improvement in accuracy, but as the resolution decreases, PINNs show a clear advantage with much smaller reconstruction uncertainty. The applicability of PINNs to empirically obtained data with no clear knowledge of the boundary condition was also demonstrated.

## Acknowledgements

The authors would like to thank S. Cai and G. Karniadakis for providing an initial framework of the PINN code and R. Thandiackal and G. Lauder for providing the PIV data for the turning zebrafish test case.

## Data availability

The PINN codes can be accessed at https://github.com/JHU-NI-LAB/PINNS_for_undulating_bodies.

## References

Agarwal, K., Ram, O., Wang, J., l, Y., and Katz, J. (2021). Reconstructing velocity and pressure from noisy sparse particle tracks using constrained cost minimization. Exp Fluids. 62, 75.

Akhtar, I., Mittal, R., Lauder, G.V., Drucker, E. (2007). Hydrodynamics of a biologically inspired tandem flapping foil configuration Theor. Comput. Fluid Dyn.. 21, 155–170.

Anderson, A., Bohr, T., Schnipper, T., and Walther, J.H. (2017). Wake structure and thrust generation of a flapping foil in two-dimensional flow J. Fluid Mech. 812, R4.

Ashraf, I, Bradshaw, H., Ha, T., Halloy, J., Godoy-Diana, R., and Thiria, B. (2017). Simple phalanx pattern leads to energy saving in cohesive fish schooling. PNAS. 114, 9599–9604.

Baydin, A.G., Pearlmutter, B.A., Radul, A.A., and Siskind, J.M. (2018). Automatic Differentiation in Machine Learning: a Survey J Mach Learn Res. 18, 1–43.

Cai, S., Wang, Z., Fuest, F., Jeon, Y.J, Gray, C., and Karniadakis, G.E. (2021). Flow over an espresso cup: inferring 3-D velocity and pressure fields from tomographic background oriented Schlieren via physics-informed neural networks J Fluid Mech. 915, A102.

Cai, S., Mao, Z., Wang, Z., Yin, M., and Karniadakis, G.E. (2022). Physics-informed neural networks (PINNs) for fluid mechanics: a review Acta Mech. Sin. 37, 1727–1738.

Cai, Z., Liu Y., Chen, T., and Liu T. (2020). Variational method for determining pressure from velocity in two dimensions Exp in Fluids. 61, 118.

Charonko, J.J., King, C.V., Smith, B.L., and Vlachos, P.P. (2010). Assessment of pressure field calculations from particle image velocimetry measurements. Meas Sci Tech. 21, 105401.

Costello J., Colin S., Dabiri J.O., Gemmell B., Lucas K., and Sutherland K.R. (2021). The hydrodynamics of jellyfish swimming. Annu. Rev. Mar. Sci. 13, 375–396.

Dabiri, J. O., Bose, S., Gemmell, B.J., Colin, S.P., and Costello, J. H. (2014). An algorithm to estimate unsteady and quasi-steady pressure fields from velocity field measurements. J. Exp. Biol. 217, 331–336.

Dabiri, J.O., Colin, S.P., Gemmell, B.J., Lucas, K.N. and Leftwich, M.C., and Costello, J.H. (2020). Jellyfish and Fish Solve the Challenges of Turning Dynamics Similarly to Achieve High Maneuverability. Fluids. 5, 2311–5521.

Dagenais, P. and Aegerter, C.M. (2020). How shape and flapping rate affect the distribution of fluid forces on flexible hydrofoils. J. Fluid Mech. 901, A1.

Dagenais P. and Aegerter C.M. (2021). Hydrodynamic stress maps on the surface of a flexible fin-like foil. PLoS ONE. 16, e0244674.

de Kat R. and van Oudheusden, B.W. (2012). Instantaneous planar pressure determination from PIV in turbulent flow. Exp Fluids. 52, 1089–1106.

Di Leoni, P.C,, Agarwal, K., Zaki, T., Meneveau, C., and Katz, J. (2022) Reconstructing velocity and pressure from sparse noisy particle tracks using Physics-Informed Neural Networks. arXiv preprint, 2210.04849 (2022).

Du, Yifan, Wang, M., Zaki, T. (2023). State estimation in minimal turbulent channel flow: A comparative study of 4DVar and PINN. Int J. Heat Fluid Fl, 99, 109073.

Fujisawa, N., Tanahashi, S., and Srinivas, K. (2005). Evaluation of pressure field and fluid forces on a circular cylinder with and without rotational oscillation using velocity data from PIV measurement. Meas Sci Tech. 16, 989.

Gemmell B.J., Du Clos K.T., Colin S.P., Sutherland K.R., and Costello J.H. (2021). The most efficient metazoan swimmer creates a ‘virtual wall’ to enhance performance. Proc Biol Sci. 288, 20202494.

Guo C., Kuai Y., Han Y., Xu P., Fan Y., and Yu C. (2022). Hydrodynamic analysis of propulsion process of zebrafish. Phys Fluids. 34, 021910.

Halsey, L.G, Wright, S., Racz, A., Metcalfe, J.D., and Killen, S.S. (2018). How does school size affect tail beat frequency in turbulent water? Comparative Biochemistry and Physiology, Part A. 218, 63–69.

Han, T., Wang, F., Wang. H., Gao, Q., and Wei, R. (2022). Experimental study on wake flows of a live fish with time-resolved tomographic PIV and pressure reconstruction. Exp Fluids. 63, 25.

He, C., Liu, Y., and Lian, G. (2020). Instantaneous pressure determination from unsteady velocity fields using adjoint-based sequential data assimilation Phys. Fluids. 32, 035101.

Huhn, F., Schanz, D., Gesemann, S., and Schröder, A. (2016). FFT integration of instantaneous 3D pressure gradient fields measured by Lagrangian particle tracking in turbulent flows. Exp Fluids. 57, 151.

Jeon, Y.J, Gomit, G. Earl, T., Chatellier, L., and David, L. (2018). Sequential least-square reconstruction of instantaneous pressure field around a body from TR-PIV. Exp Fluids. 59, 27.

Kasoju, V.T. and Santhanakrishnan, A. (2021). Aerodynamic interaction of bristled wing pairs in fling. Phys. Fluids. 33, 031901.

Kingma, D. P. and Ba, J. (2015). Adam: A method for stochastic optimization. arXiv preprint, 1412.6980.

Liao, J.C. (2006). The role of the lateral line and vision on body kinematics and hydrodynamic preference of rainbow trout in turbulent flow J. Exp. Biol. 209, 4077–4090.

Liao, J.C., Beal, D.N., Lauder, G.V., and Triantafyllou, M.S. (2003). The Kármán gait: novel body kinematics of rainbow trout swimming in a vortex street. J. Exp. Biol. 206, 1059–1073.

Li, L., Nagy, M., Graving, J.M., Bak-Coleman, J., Xie, G., and Couzin, I.D. (2020). Vortex phase matching as a strategy for schooling in robots and in fish. Nature Communications. 11, 5408.

Liu, X. and Katz, J. (2006). Instantaneous pressure and material acceleration measurements using a four-exposure PIV system. Exp Fluids. 41, 227–240.

Liu, X. and Katz, J. (2013). Vortex-corner interactions in a cavity shear layer elucidated by time-resolved measurements of the pressure field. J. Fluid Mech. 728, 417–457.

Liu, X. and Moreto, J.R. (2021). Pressure reconstruction of a planar turbulent flow field within a multiply connected domain with arbitrary boundary shapes Phys. Fluids. 33, 101703.

McClure, J. and Yarusevych. S (2017). Optimization of planar PIV-based pressure estimates in laminar and turbulent wakes. Exp Fluids. 58, 62.

McHenry, M.J., Feitl, K.E., Strother, J.A., and Van Trump, W. J. (2009). Larval zebrafish rapidly sense the water flow of a predator’s strike Biol. Lett. 5, 477–479

Mittal, R., Dong, H., Bozkurttas, M., Najjar, F.M., Vargas, A. and Von Loebbecke, A. (2008). A versatile sharp interface immersed boundary method for incompressible flows with complex boundaries. Journal of computational physics, 227, 4825–4852

Neeteson, N.J. and Rival, D.E. (2015). Pressure-field extraction on unstructured flow data using a Voronoi tessellation-based networking algorithm: a proof-of-principle study. Exp Fluids. 56, 44.

Pan, Z., Whitehead, J., Thomson, S., and Truscott, T. (2020). Error propagation dynamics of PIV-based pressure field calculations: How well does the pressure Poisson solver perform inherently? Meas Sci Tech. 27, 084012.

Pirnia, A, Mcclure, J., Peterson, S.D., Helenbrook, B.T., and Erath, B.D. (2020). Estimating pressure fields from planar velocity data around immersed bodies; a finite element approach. Exp in Fluids. 61, 55.

Raissi, M., Yazdani, A., and Karniadakis, G.E. (2020). Hidden fluid mechanics: Learning velocity and pressure fields from flow visualizations. Science. 367, 1026–1030.

Ristroph, L. and Zhang, J. (2008). Anomalous Hydrodynamic Drafting of Interacting Flapping Flags. PRL. 101, 194502.

Ristroph, L., Liao, J.C, and Zhang, J. (2015). Lateral Line Layout Correlates with the Differential Hydrodynamic Pressure on Swimming Fish. PRL. 114, 018102.

Seo, J. and Mittal, R. (2022). Improved swimming performance in schooling fish via leading-edge vortex enhancement. Bioinspiration & Biomimetics. 17, 066020

Saadat, M., Berlinger, F., Sheshmani, A., Nagpal, R., Lauder, G.V., and Haj-Hariri, H. (2015). Hydrodynamics of in-line schooling Bioinspir. Biomim. 16, 046002.

Shams, A., Jalalisendi, M., and Porfiri, M. (2015). Experiments on the water entry of asymmetric wedges using particle image velocimetry Phys Fluids. 27, 027103.

Siala, F.F, Fard, K.K, and Liburdy, J.A. (2020). Experimental study of inertia-based passive flexibility of a heaving and pitching airfoil operating in the energy harvesting regime. Phys Fluids. 32, 017101.

Tack, N.B., Du Clos, K.T., and Gemmell, B.J. (2021). Anguilliform Locomotion across a Natural Range of Swimming Speeds. Fluids. 6, 2311–5521.

Thandiackal, R. and Lauder, G.V. (2020). How zebrafish turn: analysis of pressure force dynamics and mechanical work. J. Exp. Biol. 223, 0022–0949.

Thandiackal, R., White, C.H., Bart-Smith, H., and Lauder, G.V. (2021). Tuna robotics: hydrodynamics of rapid linear accelerations. Proc Biol Sci. 288, 20202726.

van Oudheusden, B.W. (2013). PIV-based pressure measurement. Meas. Sci. Technol. 24, 032001.

Verma, S., Novati, G., and Koumoutsakos, P. (2018). Efficient collective swimming by harnessing vortices through deep reinforcement learning PNAS. 115, 5849–5854.

Wang, C.Y, Gao, Q., Wei, R.J., Li, T., and Wang, J.J. (2017). Spectral decomposition-based fast pressure integration. Exp Fluids. 58, 84.

Videler, J.J. and Hess, F (1984). Fast continuous swimming of two pelagic predators, saithe (Pollachius virens) and mackerel (Scomber scombrus): a kinematic analysis J Exp Biol. 109, 209–228.

Wang, H., Gao, Q., Wang, S., Li, Y., Wang, Z., and Wang, J. (2018). Error reduction for time-resolved PIV data based on Navier–Stokes equations Exp Fluids. 59, 149.

Wang, J., Zhang, C., and Katz, J. (2019). GPU-based, parallel-line, omni-directional integration of measured pressure gradient field to obtain the 3D pressure distribution. Exp Fluids. 60, 58.

Zhang, J., Bhattacharya, S., and Vlachos P.P. (2020). Using uncertainty to improve pressure field reconstruction from PIV/PTV flow measurements Exp in Fluids. 61, 131.

Zhang, C., Hedrick, T.L., and Mittal, R. (2015). Centripetal Acceleration Reaction: An Effective and Robust Mechanism for Flapping Flight in Insects PLoS ONE. 10, e0132093.

