## Supplementary Material for "Reconstructing the pressure field around a swimming fish using a physics-informed neural network"

### Effects of network size and training duration

The accuracy of the neural network's predictions depends on the size of the network. To determine the size of the network that generates the most accurate solution, the number of hidden layers ($L$) and the number of neurons per layer ($N$) were varied. The DNS dataset with a spatial resolution of $\Delta x = 0.02L$ and a temporal resolution of $\Delta t = 0.02T$ was used for this study. For each network size tested, the relative global root mean square error (RMSE) in the predicted velocity and pressure fields were computed as follows:

$$\text{Rel RMSE} = \frac{\sqrt{\frac{1}{N}\left[\sum_i^N \left\{(X)_{pred}^i - (X)_{DNS}^i\right\}^2\right]}}{\sqrt{\frac{1}{N_r}\left[\sum_i^{N_r} \left\{(X)_{DNS}^i\right\}^2\right]}}, \tag{S1}$$

where, $X$ is $u, v,$ or $C_p$, $N$ is the number of spatio-temporal data points in the dataset, and $N_r$ is the number of spatio-temporal data points used for the normalization. Since far away from the fish's body, the $v$ component of the velocity and the $C_p$ are both approximately zero, the normalization of these two quantifies only utilized the values near the fish's body. In this analysis, the weights of the loss function were all set to unity and the network was trained for 1000 epochs, or 155,000 iterations. One epoch is defined as the number of iterations required to make one full pass through the entire data set. As shown in Fig. S1-A, as the network size increases, the relative global RMSE decreases and eventually begins to oscillate around a mean value. Based on these results, a network size of 12 hidden layers and 120 neurons per layer was selected since it produced the smallest error in the least amount of time.

In addition to the size of the network, the duration of training also effects the accuracy of the predicted velocity and pressure field. Thus, one must determine the number of iterations that balances the computational expense with the accuracy. To test the PINN's sensitivity to the training time, it was trained for epochs ranging from 500-1750, which for this dataset corresponds to iterations ranging from 77K – 272K. For each training period, the relative global root mean square error (RMSE) in the velocity and pressure fields was computed. As shown in Fig. S1-B, 1500 epochs produces the smallest error in the pressure field predictions and requires a computational cost of roughly 9 hours. Since every 500 epochs adds approximately 1.5 hours to the computational expense, it was determined that training the PINN for 1500 epochs would efficiently balance the accuracy of the PINN predictions with the computational expense.

### Effect of weights and data noise

To test the sensitivity of the accuracy of the PINN predictions to the weighting coefficients utilized in the loss function, a weight analysis was performed where the data and boundary conditions loss terms were weighted over the Navier-Stokes equations. The weights tested included 1, 10, 50, 100, 500, and 1000. Since knowledge of the velocity field and boundary condition is needed by the PINN to find a proper solution to the Navier-Stokes equations, the idea was that by providing these terms with a higher weight, the accuracy of the method would improve. As was done in the network size and training duration studies, the relative global RMSE in the velocity and pressure fields were computed for each weight tested. In this study, the PINN was trained for 1500 epochs. As shown in Fig. S2, the error decreases significantly as the weighting coefficients approach 100. As the weighting coefficient approaches 1000, the error in the velocity field continues to decrease, but the errors in the pressure field begin to increase. This suggests that a weight that is too large would result in over fitting to the velocity field. Therefore, a weight of 100 was applied to the data and boundary condition loss terms for all cases tested in this paper. These weighting coefficients correspond to $\lambda_1$ and $\lambda_3$ in the main text.

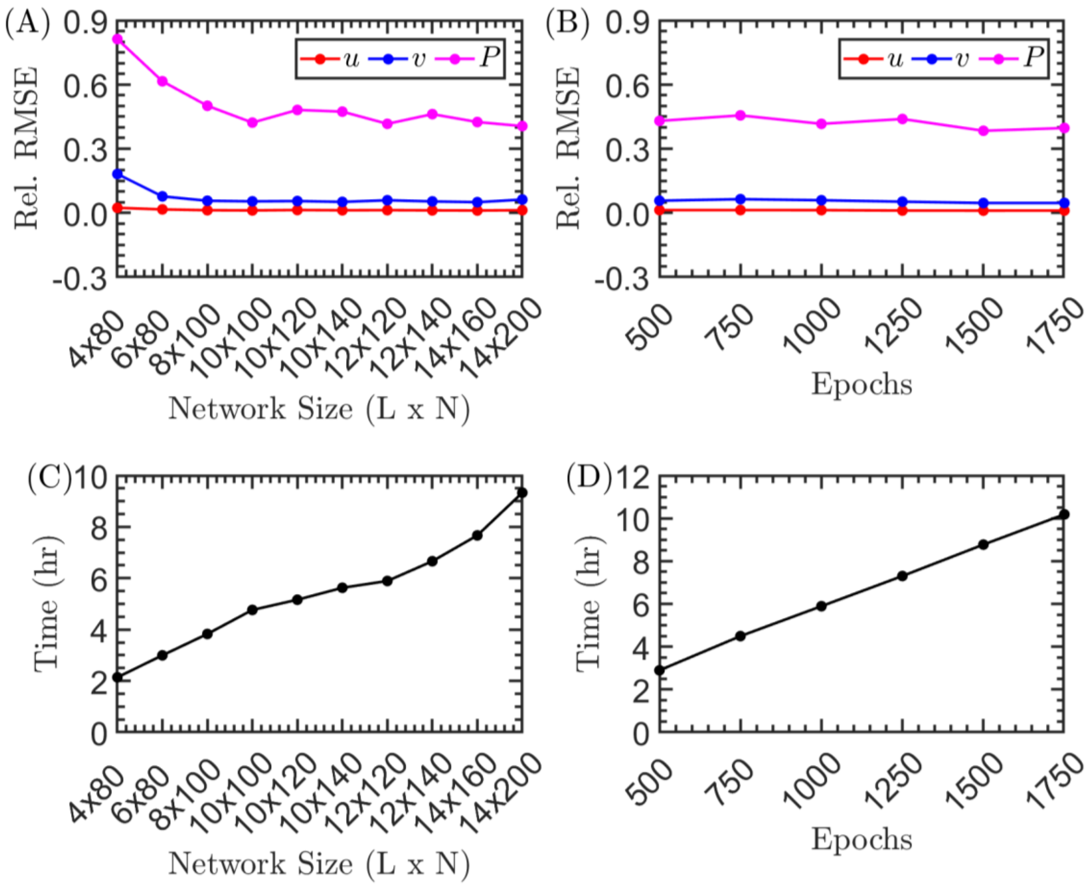


**Fig. S1. Impact of the network size and training duration on the progression of the relative global RMSE in the velocity and pressure field predictions.** Here, the network size is defined by the number of hidden layers ($L$) and the number of neurons per layer ($N$). (A) the relative global RMSE in the PINN predictions versus the network size. (B) the relative global RMSE versus the number of epochs used during the training process. The computational cost as a function of (C) network size and (D) number of epochs is also shown.

To test the sensitivity of the proposed method to noise, artificial white noise was added to the velocity data obtained on the 2D plane extracted from the DNS data. For this study, the velocity field had a spatial resolution of $\Delta x = 0.02L$ and a temporal resolution of $\Delta t = 0.02T$. Gaussian noise is considered, where the noisy velocity data is of the form:

$$u = u + a \cdot \text{std}(u) \cdot \eta \tag{S2}$$

$$v = v + a \cdot \text{std}(v) \cdot \eta \tag{S3}$$

Here, $\eta$ is Gaussian noise with zero mean and unit variance, and $a$ is the noise level defined by the ratio of the noise magnitude to the standard deviation of the velocity data. The noise level was varied from 0 to 1 by increments of 0.2. The relative global RMSE of the velocity and pressure was then computed for each noise level tested. The results are shown in Fig. S2E, which demonstrates that the PINN method is readily insensitive to noise. The errors in the pressure field only really begin to rise after a noise level greater than 0.6.

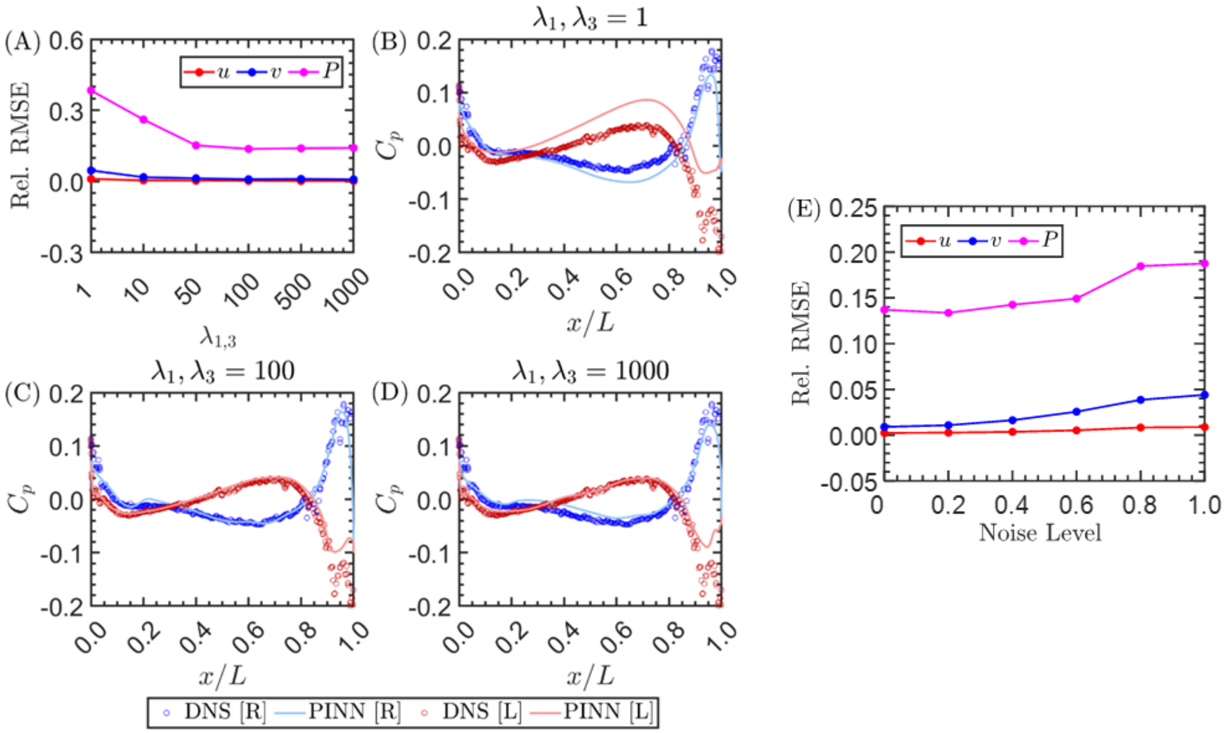


**Fig. S2. Impact of weighting the terms of the loss function and noise level on the relative global RMSE in the velocity and pressure field predictions.** (A) the global relative RMSE in the PINN predictions versus the weighting coefficient. (B)-(D) the surface pressure prediction on the left (red) and right (blue) side of the fish's body when different weighting coefficients are used. (E) Impact of the noise level on the relative global RMSE in the velocity and pressure field predictions.

### Out-of-plane Effects

As was reported in the main paper, the PINN and Queen 2.0 algorithm exhibited increased error in the tail and wake region, where the out-of-plane velocities were non-negligible. This is unsurprising since only the two-dimensional velocity field was used to reconstruct the pressure field. Thus, even if the residuals of the momentum equations shown in Eqn 8-10 of the main text are identically zero, there could still exist a certain level of error in the pressure reconstruction. This is because the residuals used in this case assumes that the product of the out-of-plane velocity and the spatial derivative of $\boldsymbol{u}$ in that direction is negligible, which is not always true. In addition, because of the missing third component from the 2D PIV, the divergence free condition was not enforced and can be used as another way to identify regions with large errors. This is demonstrated in Fig. S3, where A,C,E show the absolute error in the reconstructed pressure field at three different times in one tail beat cycle, and B,D,F show the residual of the two-dimensional continuity equation at those times. Larger errors in the pressure reconstruction occur near the body where the largest residuals of the continuity equation are found.

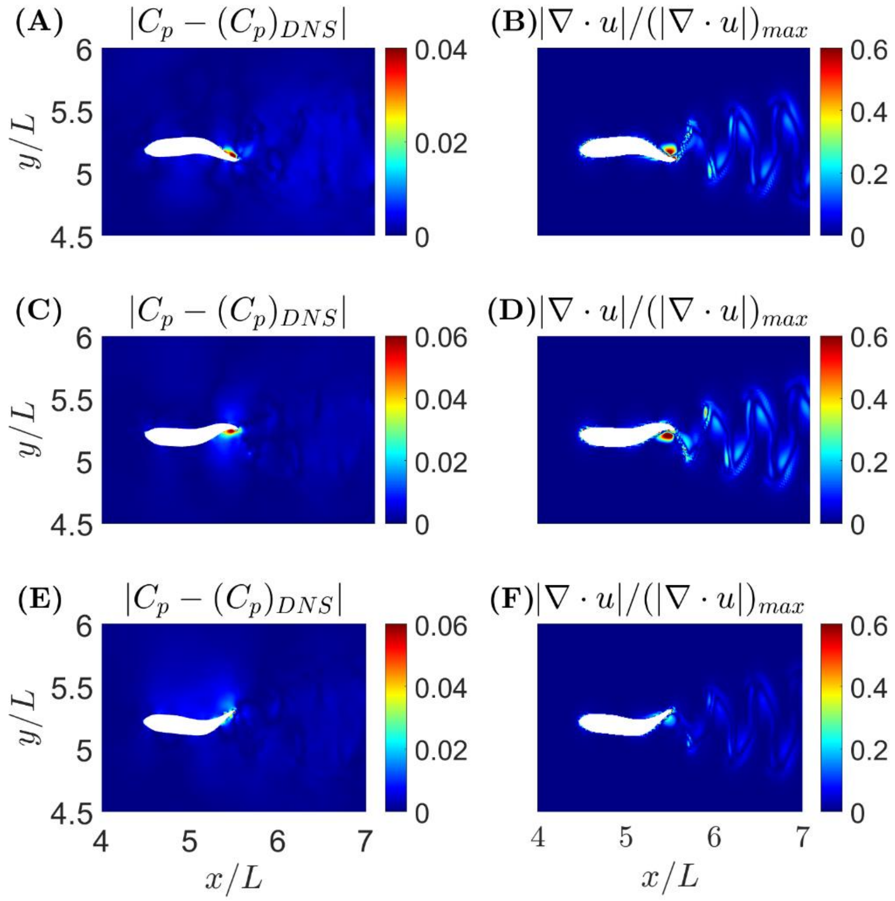


**Fig. S3. Comparison of the absolute error in the PINN prediction and the residuals of the two-dimensional continuity equation demonstrating how in areas of highly three-dimensional flow there exists an increase in pressure uncertainty.** (A)-(C) absolute error in the PINN pressure prediction at three different times. (D)-(F) corresponding residual of the continuity equation at those times.
